# Domain-specific Schema Reuse Supports Flexible Learning to Learn in Primate Brain

**DOI:** 10.1101/2024.10.27.620463

**Authors:** Kaixi Tian, Zhiping Zhao, Yang Chen, Ningling Ge, Shenghao Cao, Xinyong Han, Jianwen Gu, Shan Yu

## Abstract

Prior knowledge accelerates subsequent learning of similarly structured problems - a phenomenon termed “learning to learn” - by forming and reusing generalizable neural representations, i.e., the schemas. However, the stability-plasticity dilemma, i.e., how to exploit stable schemas to facilitate learning while remaining flexible towards possible changes, is not well understood. We hypothesize that restricting schemas to specific functional, e.g., decision-making, subspace and making it orthogonal to other subspaces allows the brain to balance stability and plasticity. To test it, we trained three macaques on visuomotor mapping tasks and recorded neural activity in the dorsolateral premotor cortex. By delineating decision and stimulus subspaces, we identified a schema-like manifold within only the decision subspace. The reuse of decision schemas significantly facilitated subsequent learning. In addition, the decision subspace exhibited a trend to be orthogonal to the stimulus subspace, minimizing interference between these two domains. Our results revealed that restricting schemas to specific functional domains can preserve useful knowledge while maintaining orthogonality with other subspaces, allowing for flexible adaptation to new environments, thereby resolving the stability-plasticity dilemma. This finding provides new insights into the mechanisms underlying brain’s capability to learn both fast and flexibly, which can also inspire more efficient learning algorithms for artificial intelligence systems towards working in open, dynamic environments.

## 1. Main

Prior knowledge can expedite the learning process, enabling new problems to be learned more quickly if they are similar to the ones we already know how to solve. This process is termed “learning to learn” [1, 24]. It is hypothesized that acquired knowledge is represented by specific neural activity patterns, referred to as schemas, which can be reused to facilitate subsequent learning [2,3,4]. Schema-like neural representations have been observed in rodents [5,6], non-human primates [7,8] and humans [9,10], manifesting as stable activity patterns across similarly structured tasks. In addition, schema formation has been observed in parallel with improved learning efficiency when mice engaged in performing sequence tasks [11]. However, regarding the scheme-based hypothesis of learning to learn, the stability-plasticity dilemma remains unsolved. That is, how to maintain stable representations so that essential knowledge can be preserved while still leaving enough plasticity to adapt to possible changes in subsequent learning (Fig. 1a).

**Fig. 1.**
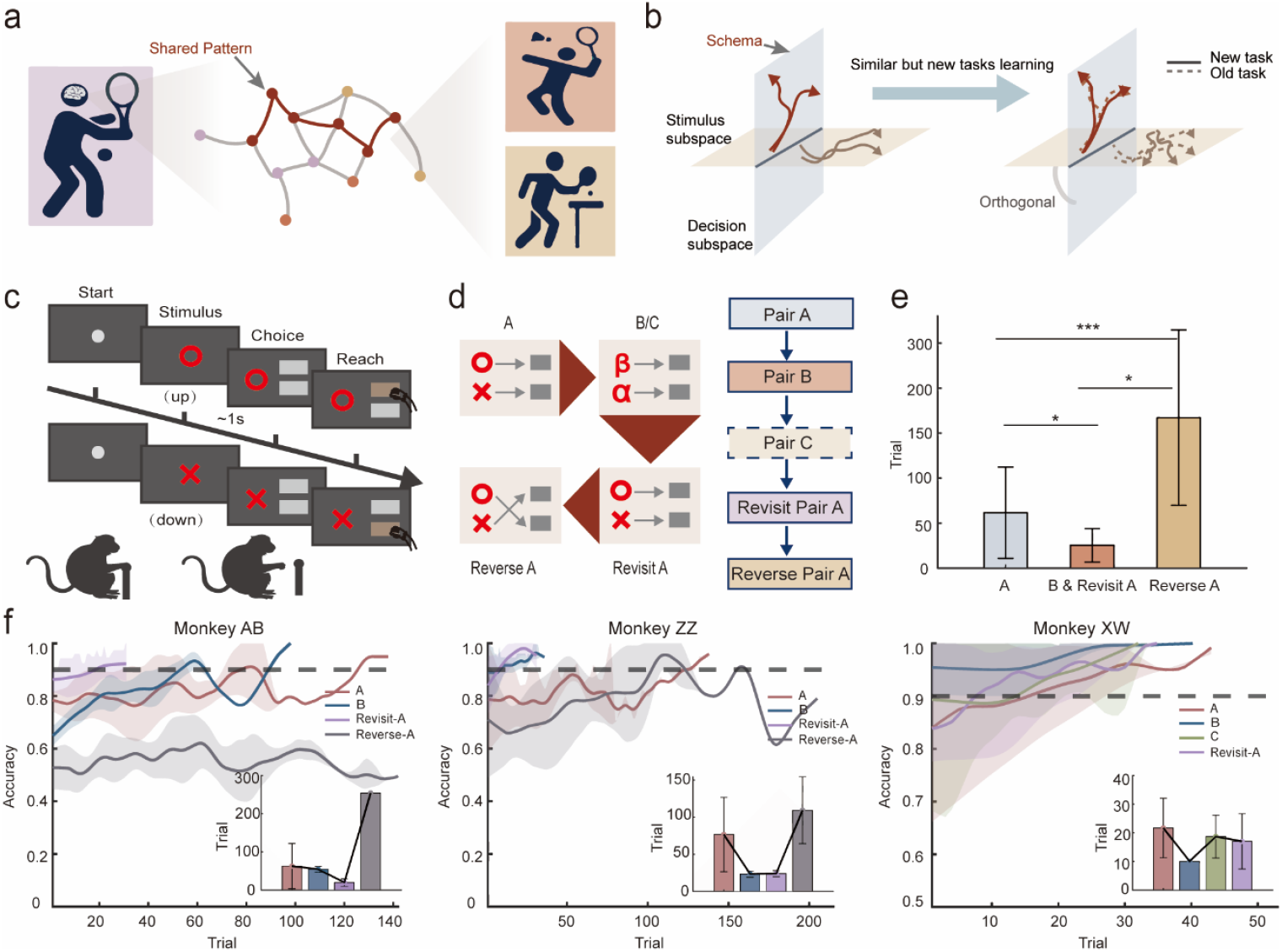
Schematic of problems and hypotheses, experimental design and behavioural analysis. (a) Illustration of the balance between schema reuse and flexibility, e.g., individuals skilled in tennis can more rapidly acquire related skills in other racquet sports, such as badminton and table tennis, despite significant changes across individual sports. (b) Hypothesis: Schema reuse is preserved in a specific domain to keep the essential knowledge or skill, while other functional domains that are orthogonal to it can adapt to changes. (c) Schematic of the visuomotor mapping tasks. (d) Workflow for learning a series of visuomotor mapping tasks. (e) Statistics on the number of trials required by two monkeys (monkeys AB and ZZ) to learn various tasks. (f) Learning speed across tasks for three monkeys, with the grey dashed line indicating the criterion for task mastery. The shaded area represents the variance of accuracy across different days. The inset shows the number of trials required to reach that criterion. \**P* < 0.05, \*\**P* < 0.01, \*\*\**P* < 0.001.

The stability-plasticity dilemma presents a significant challenge in the field of machine learning, regarding how intelligent systems can leverage prior tasks or experiences to inform future learning. Various approaches have been employed to address this issue: continual learning explores the problem of catastrophic forgetting in artificial neural networks [12,13,14], while meta-learning focuses on how to rapidly adapt to new tasks through a few trials [15,16], and online learning emphasizes strategies for tackling unknown tasks in dynamic environments [17]. Despite these advances, it is still challenging for artificial intelligence systems to balance stability—retaining past knowledge while minimizing catastrophic forgetting—and plasticity, which involves the ability to learn quickly from new experiences and generalize effectively. In contrast, biological systems, particularly mammals, exhibit remarkable balance between stability and plasticity: they preserve and effectively capitalize on prior knowledge, while still being highly flexible in dealing with unforeseeable changes [18,19].

Recently, Goudar et al. used an artificial neural network model to simulate the learning of a series of visuomotor mapping tasks [20]. The model exhibited a schema-like, low-dimensional activity manifold, accompanied by increased learning efficiency. The authors demonstrated that the schema and its reuse were observed only in the decision subspace, but not in the stimulus subspace. Similarly, Laura et al. achieved flexible multi-task computation in artificial neural networks through reusing of local dynamical motifs [21]. These studies inspired us to hypothesize that restricting the schema to specific functional domains, e.g., decision, and making it orthogonal to other subspaces allows the brain to preserve essential knowledge while enabling flexibility to handle the variability of future problems, e.g., with different stimuli, thereby providing a solution to the stability-plasticity dilemma (Fig. 1b).

To test this hypothesis, we trained three macaques on a series of visuomotor mapping tasks and recorded neural population activity in the dorsolateral premotor cortex (PMd) (Supplementary Fig. 1), a region involved in decision-making and action planning [22, 23]. We delineated the decision and stimulus subspaces to search for the schema and examine how its reuse can facilitate learning. The tasks required the monkeys to press corresponding buttons based on the presented visual stimuli (Fig. 1c). Each day, the monkeys needed to learn two (task A and B for monkeys AB and ZZ,) or three (task A, B and C for monkey XW) new stimulus-action pairs, followed by a task of revisiting pair A (Revisit-A). For monkeys AB and ZZ, an extra reverse task was added at the end, in which the animals needed to learn the reverse mapping of pair A (Reverse-A) (Fig. 1d). Data from three days for each monkey were analysed (monkey AB/ZZ/XW completed 257±27/212±41/209±7 trials per day). In total, we recorded activities from 948 multi-unit (MUA) and single-unit (SUA). Only the activities during the 1-second visual cue presentation time, before the onset of the action, were analysed in the present study (see Methods for details).

The behavioural results are shown in Figure 1e and 1f. Similar to previous findings [11, 24], we found that learning efficiency (measured by the number of trials needed to reach a 0.9 correct rate in 20 consecutive trails) increased for later-learned pairs. Examining each monkey individually (Fig. 1f) shows that the learning speed for pairs B/C was faster compared to the pair A in monkeys ZZ and XW, and the learning speed for the revisit-A was faster compared to the pair A in two monkeys AB and ZZ. In contrast, the reverse task was more challenging, with the learning speed of Reverse-A significantly slower than that of A, B/C, and Revisit-A (Fig. 1f). Pooling the data from two monkeys (AB, ZZ) that finished learning pair A, B, Revisit-A, and Reverse-A together reconfirmed the trends we saw in individual animals, that is, later learned pairs exhibited significantly increased efficiency, while the reverse learning task was significantly slowed (Fig. 1e). The analysis of the reaction time (RT) also corroborated that of the accuracy (Supplementary Fig. 2). Overall, these behavioural results reconfirm that similar problems encountered later can be learned faster, but learning new problems may be delayed if they contradict previously acquired knowledge (see supplementary discussion 1.1).

**Fig. 2.**
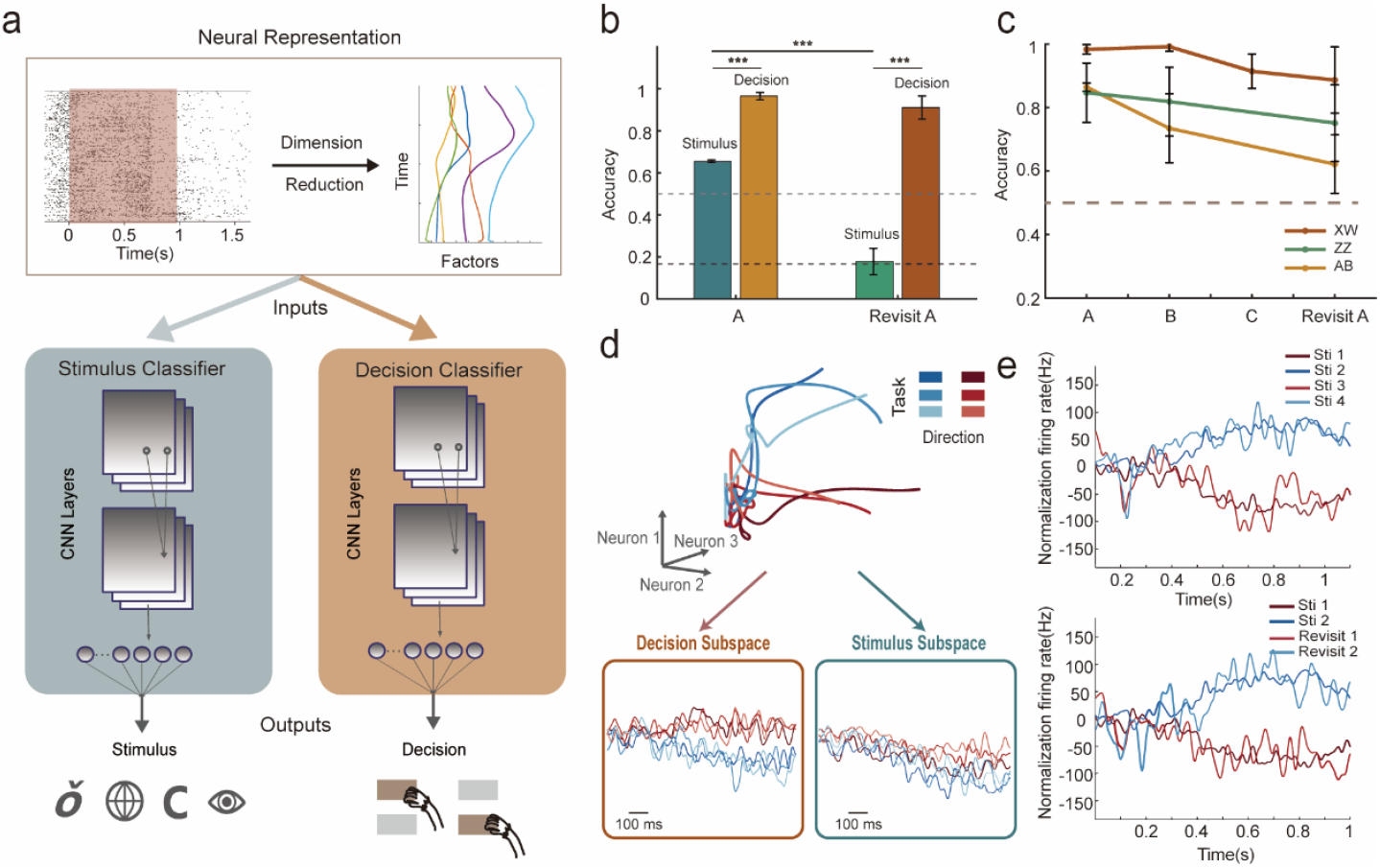
Identifying shared patterns from neural population activity. (a) Schematic of the decoding shared patterns across tasks: neural population activity from task A was reduced to a lower dimension using a nonlinear method, and CNNs were trained separately to classify different visual stimuli and motor decisions. The reddish shadow area highlights neural activity from the onset of visual stimulus presentation to the appearance of the choice button. (b) The trained classifiers were used to decode visual stimuli and motor decisions, with classification accuracy computed for task A and generalized to the task of Revisit-A. The dark gray dashed line represents the chance level for visual stimulus classification, and the light gray dashed line represents the chance level for decision-making classification. (c) The accuracy of the decision classifier for monkey XW trained in task A was applied to tasks B/C/Revisit-A. The gray line represents the chance level of decision classification. (d) Decompose neural population activity into the decision subspace and the stimulus subspace for monkey AB. (e) Neural activity during subsequent learning tasks from monkey XW was projected into the decision subspace using the projection matrix from task A. The top panel shows task B, and the bottom panel shows Revisit-A. Sti, stimulus; XW, monkey XW; ZZ, monkey ZZ; AB, monkey AB; \**P* < 0.05, \*\**P* < 0.01, \*\*\**P* < 0.001.

Next, we searched for schemas in population activities recorded from the PMd, decomposed to the stimulus and decision domains. First, we examined whether the population activity indeed encoded task-related parameters in the visual and decision domains. To this end, we constructed two classifiers using convolutional neural networks (CNNs) to decode visual stimuli and motor decisions (Fig. 2a). We employed a nonlinear dimensionality reduction method (LFADS) [13] to reduce the high-dimensional neural firing rate data to a low-dimensional space. The data recorded during the learning of new pairs were used to train the classifiers. We found that PMd activities encoded both visual stimuli and motor decisions, with the corresponding classifiers exhibiting significantly higher performance compared to the chance level. We note that the encoding of visual stimuli was independent of the motor decisions, as the visual classifier could discriminate different stimuli associated with the same motor response (Fig. 2b). Results for monkey XW are shown on the left side of Figure 2b, and similar results for monkeys AB and ZZ are shown in Supplementary Figure 3a. To test whether these information-bearing neural activities were stable across tasks, we fixed the parameters of the two classifiers and calculated their performance in the Revisit-A task. Surprisingly, even with the same stimuli as in task A, the visual classifier failed to discriminate the visual stimuli, exhibiting only chance level performance (the right side of Fig. 2b, monkey AB, ZZ’s results in Supplementary Fig. 3a), suggesting that the visual representation in the PMd was not stable across tasks. In contrast, the motor decision classifier exhibited consistent performance when applied to the unseen data from Revisit-A, compared to the data on which it was trained (the right side of Fig. 2b, monkey AB, ZZ’s results in Supplementary Fig. 3a), suggesting the decision-related information was stable across tasks.

To further explore the across-tasks stability of decision-related representations, we trained another motor decision classifier using data only from task A and tested its generalization ability on tasks B/C. We found that the classifier trained on task A data successfully generalized to the unseen data from tasks B/C and Revisit-A (Fig. 2c), confirming that in similar tasks, the decision-related representation was largely shared across individual tasks.

The classifier analyses demonstrated the existence of schema-like, stable representations in the domain of motor decisions. Next, we examined more explicitly how such representations are embedded in population dynamics. The results of monkey AB are shown in Figure 2d, with similar results found in the other two monkeys (Supplementary Fig. 3b). We used demixed principal component analysis (dPCA) [14] to project population activity collected in tasks A, B/C and Revisit-A into the stimulus and decision subspaces. In the decision subspace, neural dynamics associated with the same decision across different tasks were clustered and separated from the cluster representing the opposite decision (Fig. 2d, lower-left), suggesting that the schema was represented as dynamics within a low-dimensional manifold in the decision subspace. Consistent with the classifier results, the population dynamics representing different stimuli were largely mixed (Fig. 2d, lower-right).

We further examined the reuse of low-dimensional manifolds in the decision subspace across individual tasks. First, we conducted the dimension reduction for task A data in the decision subspace. Then, we projected the data from tasks B/C and Revisit-A into the same subspace. We found that, similar to the results described above, the decision manifolds obtained from task A were reused for other, similar tasks (Fig. 2e, monkey AB, ZZ’s results in Supplementary Fig. 4).

Next, we analyzed the population activities in the Reverse-A task. It helps to discriminate if the reused neural manifolds indeed reflect the decision *per se*, or merely reflect the planned movement direction. If the latter was the case, we would expect that the reuse can be found in the Reverse-A task as well, since the movement directions in this task were the same as in other tasks. The results show that the opposite was true, i.e., stable neural manifolds shared across tasks A, B/C, and Revisit-A were not reused in the Reverse-A task (Fig. 3a, 3d). Quantitative analysis confirmed that the similarity of the manifold in the decision subspace between the task A and Reverse-A was significantly lower than that between the task A and tasks B/C/Revisit-A (Fig. 3b, 3e), suggesting that the information encoded in these representations was decision-related, rather than movement-related.

**Fig. 3.**
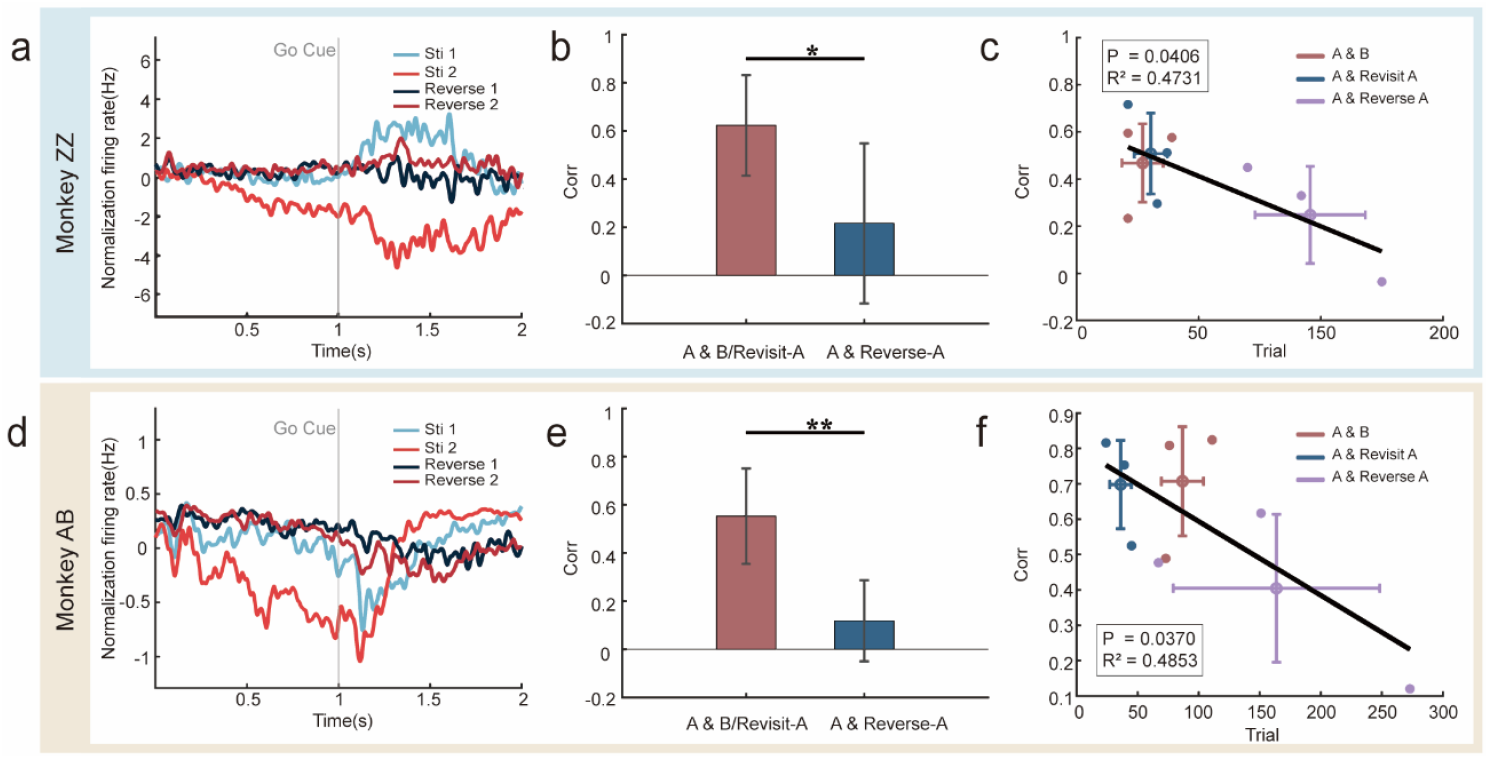
The decision representations in the reverse task. (a,d) The neural manifold of the decision subspace between task A and task Reverse-A. (b,e) The similarity between the representation of the decision subspace manifold in A/B/Revisit-A tasks and in the Reverse-A task. (c,f) A linear relationship exists between the similarity of the representation of the decision subspace manifold and the number of trials required to learn the tasks.

Then, we examined the functional role of the schema reuse in the decision subspace identified above. In Figure 3c, 3f, for monkeys AB and ZZ, we plotted the number of trials needed each day to learn the task B, Revisit-A and Reverse-A versus the degree to which the schema identified in task A was reused (measured by the correlation between the manifolds in the decision subspace). The results show a significant negative relationship between the two. In the case of the Reverse-A task, learning was actually delayed, as it required more trials to learn compared to the task A. These results suggest that whether the schema can be reused may strongly affect subsequent learning, i.e., it may facilitate learning for similar tasks but delay learning for dissimilar tasks.

We have demonstrated above that the schemas were only preserved and reused in the decision but not the stimulus subspace. Next, we investigate the neural mechanisms that minimize the interference between the representation of decision and sensory stimulus. Computationally, an orthogonal relationship is the ideal form to minimize possible interference between different dimensions. To test if it is the case between the decision subspace and the visual stimulus subspace t examined in the current study, we identified the major components in these two subspaces that explained the highest variance in the data, and then calculated the angle between them (see methods for details). As a control, we used a shuffling strategy to randomize the stimulus and decision labels for each trial, recalculating the angular separation between the subspaces. We found that the angle between the actual data was significantly closer to orthogonal than the shuffling results (Fig. 4a). Consistently, the scatter plot illustrates that, along the visual stimulus dimension, different visual stimuli were well distinguished, while motor decisions were not; conversely, in the decision dimension, motor decisions were clearly differentiated, whereas visual stimuli were not (Fig. 4b, the results of monkey AB; monkey ZZ’s results are shown in Supplementary Fig. 5). To better visualize the orthogonal representation, we plotted the neural dynamics in three-dimensional space. Figure 4c illustrates the orthogonal-like relationship between these two subspaces. We selected a day with an average angle between the two subspaces of 81°for monkey AB to depict the temporal evolution of neural signals along these axes. As a control, an example with a 25°angle is also shown, which was calculated for minor components in the two subspaces that explained much less variance. Clearly, when the decision subspace is approximately orthogonal to the visual stimulus subspace, visual stimuli and motor decisions can be effectively separated without interference (Fig. 4d, left side). However, when the relation is far from orthogonal, there is significant overlap between the two subspaces, distinguishing different visual stimuli and classifying motor decisions becomes intermingled (Fig. 4d, right side). Taken together, these results indicate a near orthogonal relationship between the decision and visual stimulus dimensions, which may contribute to non-interfering representation of decision and sensory stimulus.

**Fig. 4.**
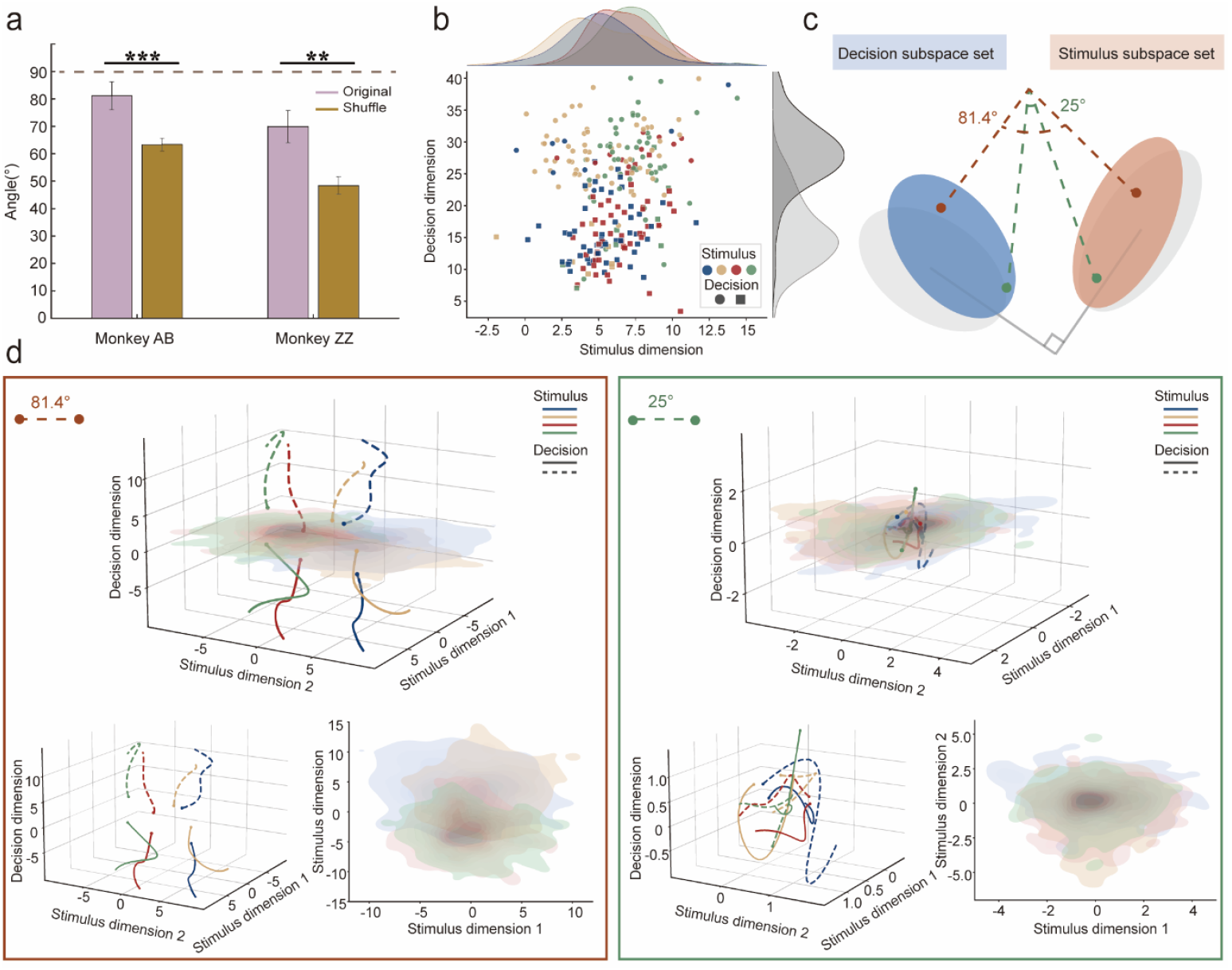
Orthogonal relationship between decision subspace and stimulus subspace. (a) The angle between the stimulus subspace and the decision subspace before and after shuffling, shown for the largest variance component. The error bar represents the variance of angles across different days. (b) Distribution of data for individual trials of monkey AB on the stimulus-decision plane; (c) Schematic illustration of the angle between the stimulus subspace and the decision subspace. (d) The left panel illustrates the neural dynamics trajectory in the three-dimensional space of the decision and stimulus subspaces when the angle is 81.4°, along with projections of motor decisions and visual stimuli. The right panel shows the neural dynamics trajectory in the three-dimensional space of the decision and stimulus subspaces when the angle is 25°. *P < 0.05, **P < 0.01, ***P < 0.001.

## 2. Discussion

In the present study, we demonstrated that schema-like, stable neural activities are embedded in the low-dimensional manifolds in the PMd of macaques during the process of learning a series of visuomotor mapping tasks. The schema was represented and reused only in the decision subspace, and its reuse strongly affects subsequent learning, providing strong evidence for the schema-based hypothesis of learning to learn [20].

Theoretically, by restricting the formation, preservation and reuse of the schema in the decision subspaces, subsequent learning of similar rules can be accomplished by “within-manifold perturbation”, which can be implemented much easier [25, 26]. At the same time, this also frees other task-related representation subspace, e.g. the subspace representing visual stimuli or the motor responses, so that they can be modified to handle various sensory inputs and motor outputs in solving new problems. Our findings, for the first time, demonstrate that this is indeed the case for the primate brain, when it is engaged in a series of tasks that require quick and flexible learning. In addition, we found that such domain-specific preservation and reuse of schemas may rely on an approximately orthogonal relationship between the decision and visual subspaces. This not only reveals the computational mechanism underlying the effective use of schemas in the brain, but also suggests a possible solution for artificial intelligence systems to preserve and reuse key knowledge in multi-task learning [27, 28]. The PMd cortex simultaneously encodes sensory evidence and decision-making behavior [29], and it can retrieve and maintain behavioral goals based on the identity of visual stimuli without encoding the visual features of objects [30]. The PMd is closely connected anatomically to the parietal cortex, likely receiving visual information through this connection [31, 32]. The PMd also receives limited input from the vlPFC and dlPFC, both of which are involved in processing visual object information [33, 34, 35]. Therefore, it is conceivable that the PMd encodes the identity features of visual stimuli. Thus, the PMd may combine visual information with behavioral goals, thereby playing a pivotal role in goal-directed motor planning and execution [36]. There is evidence that high-level cortical areas such as the motor, premotor, and prefrontal cortices have the ability to encode and integrate information across multiple dimensions, providing the neural foundation for complex and flexible cognitive functions [37,38]. Orthogonal coding is key to the parallel processing of multidimensional information [39]. Evidence of orthogonal coding representing different attributes of objects and contextual information has been found in several brain regions [40, 41]. For example, Yiling et al. identified orthogonal coding of visual and motor information in the visual cortex [42]. Zhang Y et al. discovered orthogonal coding of sensory and motor information in the motor cortex of monkeys [43]. Timo Flesch et al. indicates that projecting task representations onto low-dimensional orthogonal manifolds can convey richer information, which has been validated in the human prefrontal and posterior parietal cortices [44, 45]. These findings suggest that orthogonal coding may be a widespread form of representation in the brain, enabling parallel computation. Our finding adds to this understanding by revealing that orthogonal coding can minimize interference among various domains. This is critical for effective compartmentalization of different processing even within the same cortical region, and as the example we showed here it supports flexible use of domain-specific schemas.

Interestingly, our study also highlighted the challenges faced by monkeys in reversal learning tasks, which differ from those in similar learning situations. Reversal tasks are an important experimental paradigm for assessing cognitive flexibility [46]. The neural mechanisms underlying reversal learning tasks are not well understood. In the field of cognitive psychology, some studies have proposed an inseparable relationship between cognitive flexibility and schemas [47]. Our results showed that the lack of schema-like neural dynamics in reversal tasks suggests that learning reversal mapping rules may rely on different neural mechanisms. These differences prompt further consideration of the relationship between cognitive flexibility and schema utilization. This is consistent with previous psychological studies indicating that disrupting schemas may enhance cognitive flexibility [48].

Previous studies have found neural schemas mostly in higher cognitive brain regions such as the orbitofrontal cortex, hippocampus, and prefrontal cortex [49]. However, we also observed the reuse of schemas in the PMd, a region closely associated with cognitive flexibility and motor control. Given that schemas facilitate the reuse of prior knowledge in adaptive learning, it is plausible that this mechanism is not confined to specific brain regions, but is a fundamental organizational principle that extends across various cortical areas. The ability to preserve and adapt previously learned rules could be critical for many cognitive processes, such as decision-making, sensory processing, and motor control, implying that schema-like neural dynamics may be a general mechanism for efficiently managing cognitive flexibility and stability in a wide range of tasks.

In summary, our results offer a neural mechanism for resolving the stability-plasticity dilemma by embedding reusable schemas in specialized subspaces, ensuring both the preservation of prior knowledge and the adaptability required for new learning. This computational framework not only deepens our understanding of schema-based learning in the brain but also suggests potential strategies for artificial intelligence systems designed to balance stability and flexibility across multiple tasks. Moreover, the evidence of schema-like neural dynamics in the PMd supports the hypothesis that schemas may be a universal organizing principle across cortical regions, involved in preserving and adapting knowledge for diverse cognitive and motor functions.

## 3. Materials and Methods

### Animals and experiment conditions

Three male macaques (monkey AB weights 9.8kg, aged 9 years; monkey ZZ weights 10.6kg, aged 10 years; monkey XW weights 8.9kg, aged 6 years) were trained to perform visuomotor mapping tasks. All experimental protocols were approved by the Animal Care and Use Committee of the Institute of Automation, Chinese Academy of Sciences, in accordance with ethical guidelines. During the task, the monkey was seated in a chair, with heads fixed through a surgically implanted titanium post and left hands constrained. A 17-inch capacity touch screen (60Hz frequency; 1024×768 dpi) was placed in front of the monkey. The animals used their right hands to touch the screen to perform the tasks. A fixed grip bar with two built-in infrared sensors (EE-SPY302-1, AOYOU) was mounted on the front of the monkey chair. The infrared sensor detects whether the monkey’s hand was gripping the bar.

Monkey XW’s left PMd cortex was implanted with a 96-channel Utah array (400 um pitch, electrode length 1mm; Blackrock, USA). Monkey AB and ZZ’s left PMd cortices were implanted with two 48-channel Utah arrays (double-headed electrode with 48 channels per head, 400 um pitch, electrode length 1mm; Blackrock, USA) (Supplementary Fig. 1). A Cereplex Direct (Blackrock, USA) multi-channel data acquisition system was used to collect data in the frequency range of 0.1 Hz - 5500 Hz with a sampling rate of 30,000 Hz. The parameters in the Mountainsort [50] program were customized to automatically classify single-unit and multi-uni. Specifically, a 250 Hz Butterworth high-pass filter was used to process the raw data. A threshold of -5.5 times the standard deviation was used to detect action potentials within each electrode. When the distance between an action potential waveform and the cluster it belongs to was greater than or less than 3 times the MSE (mean squared error) of the cluster distribution, the action potential waveform was defined as background noise. A unit without an inter-spike interval (ISI) of less than 2 ms and an isolation score greater than 2.33 from other clusters was defined as a single unit. The rest were marked as multiple units. SU and MU were pooled together to form the population activities, which were analyzed in the present study.

### Behavior tasks

The monkeys were required to learn a series of new visual stimuli-action pairs, revisit a previously learned pair, and learn the reverse mapping of a previously learned pair (Fig. 1a). At the beginning of each session, a red dot appeared in the middle of the monitor. When the monkey held the bar for 1 second, the red dot disappeared, and a visual stimulus was presented on the screen for 1 second. Immediately after the offset of stimulus, two buttons appeared on the upper-right and lower-right positions of the screen as the Go Cue, prompting the monkey to release the bar and start hand moving. The monkey needed to touch the correct button according to the visual stimulus presented within 5 seconds to receive a juice reward. Incorrect trails were indicated by a specific sound followed by a black screen for 5 seconds.

### Data analysis

The behavioral and neural activity data from three monkeys over three days were analyzed. For the behavioral data, a sliding window of 20 trials was used to calculate the accuracy. Since monkeys AB and ZZ performed the same sequence of tasks, their data were combined for statistical analysis. Reaction time (RT), defined as the time it takes for the monkey to leave the bar after seeing the go cue and touch the selected button, was calculated for different tasks. To evaluate significant differences in RT between different tasks, a two-sample t-test was employed (P>0.05 to reject ; P<0.05,P>0.01*; P<0.01,P>0.001,**;P<0.001,***).

The spike data were structured into a continuous binary raw format within one second before the onset of Go cue. The spike count was binned at 10 ms intervals, resulting in a matrix, *X* ∈ ℝ^*M*×*N*×*T*^, where *M* is the number of trials, *N* is the number of neurons, and T is the number of spikes across time. The LFADS [51] method was employed to perform non-linear dimensionality reduction on the neural spike data, reducing it to 16 dimensions. The low-dimensional neural data matrix was represented as *X*_*l*_ ∈ ℝ^*M*×16×*T*^. Subsequently, a two-layer CNN was used to classify the reduced neural data in response to image stimuli and motor decisions in similar learning tasks.

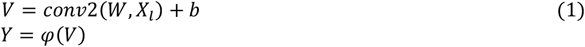

where *conv*2 is the function for 2D convolution, *W* is the weight matrix of the convolutional layer, *b* is the bias, and *ϕ* is the activation function *Relu*. The network architecture for classification consists of two convolutional layers, each with *n*_*filter* = 8, *kernel* = 3, *stride* = 1, *padding* = 1, and one fully connected layer, with *dropoutrate* = 0.5. Data from the series of new stimulus-action pair learning tasks were used to train both the visual stimulus classifier (Note: Monkey XW classified 6 categories of images, while monkeys ZZ and AB classified 4 categories) and the motor decision classifier (all three monkeys classified 2 categories, up and down). The dataset was divided into 80% for training and 20% for testing. With the stimulus model parameters *W* fixed, the unseen neural data from the revisit-A task were fed into the same CNN model to classify visual stimuli. Similarly, with the decision model parameters fixed, the neural data from the revisit-A task were fed into the same model for decision classification. To further validate, a decision classifier was trained using only the data from task A. The fixed-parameter model was then tested using neural data from task B/ C and revisit-A.

Manifolds related to motor decision-making and visual stimuli were derived from the neural activity in the PMd [52]. Spike trains for each trial were smoothed using a Gaussian kernel (σ = 50 *ms*). Neural data were represented as *X* ∈ ℝ^*N*×*S*×*D*×*M*^, where *M* is the maximum number of trials across all conditions, *N* is the number of neurons, *S* is the type of visual stimulus, and *D* is the direction of motor decision-making. Trials under different conditions were averaged to obtain *X*_*ave*_ ∈ ℝ^*N*×*S*×*D*^, which was then marginalized [53] into different factors: *X*_*ave*_ = *x*_*t*_ + *x*_*ts*_ + *x*_*td*_ + *x*_*tsd*_ + *x*_*noise*_, where *x*_*t*_ represents condition independent term, *x*_*ts*_ the stimulus term, *x*_*td*_ the decision term, *x*_*tsd*_ the stimulus-decision interaction term, and *x*_*noise*_ the noise term. Dimensionality reduction was then applied to the demixed components to extract the subspace *Y* ∈ ℝ^*p*×*S*×*D*×*T*^, where *p* is the number of principal components (PCs) used for further analysis (*p* = 20), with each dimension corresponding to a specific term noted before. The eigenvector matrix was denoted as *W* ∈ ℝ^*N*×*p*^. The decision dimension was referred to as the decision subspace 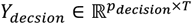, where *p*_*decision*_ is the number of decision components, and the stimulus dimension as the stimulus subspace 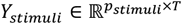, where *p*_*stimuli*_ is the number of stimulus components. For further analysis of manifold reuse, dPCA was performed on the neural data only from task A to obtain the projection matrix *W*_*A*_. Neural data from tasks B/C and revisit-A were projected into task A’s subspaces using the same matrix *W*_*A*_.

In the decision subspace *Y*_*decsion*_, the dimension with the highest proportion of variance *S* ∈ ℝ^*S*×*D*×*T*^ was selected. The Pearson correlation coefficient *rho*_*i,j*_ between the upward and downward decision manifolds in the similar learning tasks were calculated, where *cov* represents covariance, *rho* is the Spearman rank correlation coefficient, *i* refers to task A, *j* refers to task B/Revisit-A/Reverse-A, *c* indicates whether the motor decision direction was up or down, and 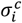 is the variance of 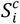. The correlation was calculated as:

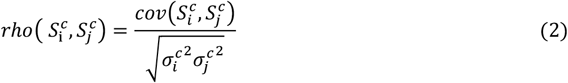

The average correlation coefficient between the two decisions was taken as the decision manifold correlation coefficient to measure the degree of manifold reuse. A two-sample t-test was conducted to assess significant differences in the decision manifold correlation coefficient between the task Reverse-A and the tasks A, B/C and Revisit-A. Linear regression analysis was performed on the reaction time and the decision manifold correlation coefficient, and the p-value and the linear regression coefficient *R*^2^ were calculated.

The angle between the visual stimulus subspace and the motor decision subspace was calculated by identifying two dimensions in the decision subspace *D*_*d*2_ that explained the highest variance and accounted for more than 1%, along with three dimensions in the stimulus subspace *D*_*v*3_. This angle is denoted as *θ, θ* = asin (min (1, *norm*(*D*_*v*3_− *D*_*d*2_× (*D*_*d*2_ × *D*_*v*3_)))). The shuffle involves randomly scrambling the labels of visual stimuli and motor decisions. Using the same angle calculation method, a one-sided t-test was conducted for significance testing. The three-dimensional neural dynamics schematic selected as an example corresponds to an angle of 81.4°between the visual stimulus subspace and decision subspace for monkey AB, while the counterexample involves an angle of 25°between dimensions of decision and visual stimuli that explain less than 1% variance for monkey AB.

## Notes

### Competing Interest Statement

The authors have declared no competing interest.

