## Supplemental materials for "Domain-specific Schema Reuse Supports Flexible Learning to Learn in Primate Brain"

**Supplementary Information**

1. **Supplementary Notes**
1.1 **Behavioural Results**

It is noteworthy that our monkeys had previously been trained on fixed stimulus pair, providing them with a foundational understanding of the task framework compared to learning a completely new task. During training, we defined task mastery as achieving an accuracy of 80% over 50 consecutive trials. In Figure 1f, the number of trials required for training is depicted by different colored lines for each monkey, with the lines for the B/C and Revisit-A tasks being notably shorter than for the A task. Particularly, monkey AB and ZZ required fewer trials for the Revisit-A task compared to the B task. Monkey XW, having already mastered the task framework, exhibited rapid learning, mastering tasks within approximately 10 trials, making differences less noticeable. Nevertheless, all three monkeys demonstrated a trend of increasingly faster learning, consistent with other behavioural studies [11, 20, 24].

For a more detailed analysis, we revised the criterion during data analysis: achieving 90% accuracy over 20 consecutive trials was considered task mastery. Based on this criterion, the data from monkey AB and ZZ (Fig. 1e) revealed that the learning rate for the Revisit-A task was significantly higher than for the A task, while the reverse task proved significantly more challenging. Individual analyses of each monkey, shown in the bar plots of Figure 1d, confirmed similar results.

Regarding reaction times across different tasks (Supplementary Fig. 2), monkey AB and ZZ exhibited significantly faster reaction times for the B and Revisit-A tasks compared to the A task, while reaction times increased significantly during the reverse task. For monkey XW, reaction times progressively slowed, possibly due to overfamiliarity with the tasks, affecting motivation and other factors, making reaction time an unreliable indicator of task mastery. Overall, the monkeys showed a trend of faster learning in the A, B/C, and Revisit-A tasks, but faced greater difficulty during the reverse task.

2. **Supplementary Figures**

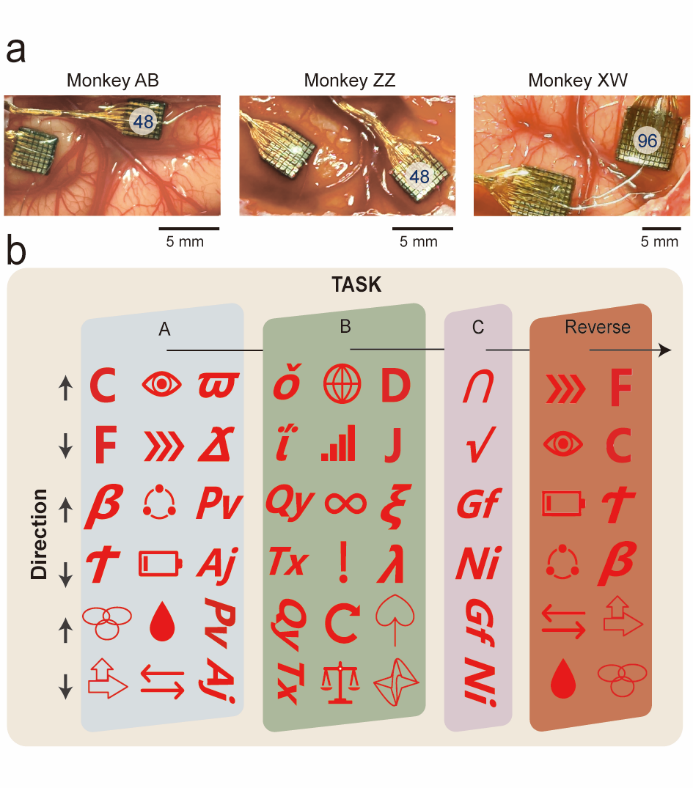

Supplementary Fig. 1. Electrode implantation and visual stimulus pairs displayed in tasks (a) Implant UTAH electrodes in three monkeys. (b) The set of Visual stimulus pairs of visuomotor mapping tasks.

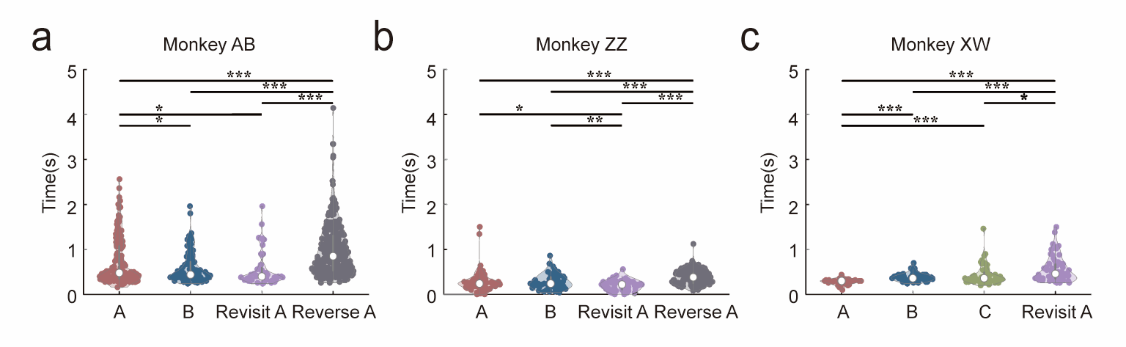

Supplementary Fig. 2. (a,b,c) Response time (The time interval from Go Cue to monkey’s hands leaving the grip) of monkeys learning different tasks. **P* < 0.05, ***P* < 0.01, ****P* < 0.001.

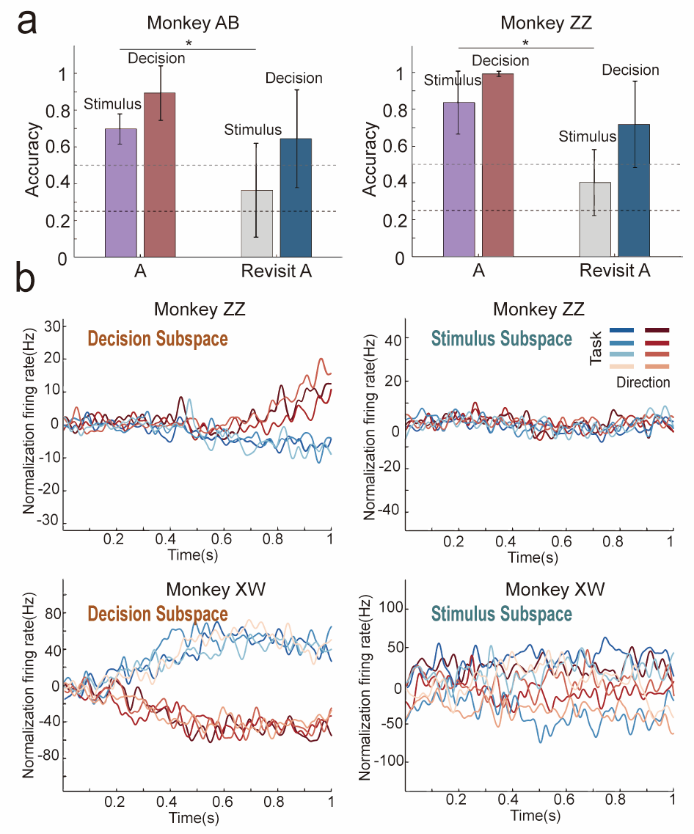

Supplementary Fig. 3. (a) Classifiers trained on Task A in two monkeys were used to decode visual stimuli and motor decisions. The classification accuracy was tested on Task A and generalized to the revisit A task. The dark gray dashed line represents the chance level for visual stimulus classification, while the light gray dashed line indicates the chance level for decision classification. **P* < 0.05, ***P* < 0.01, ****P* < 0.001. (b) Decompose neural population activity into the decision subspace and the stimulus subspace for monkey ZZ and monkey XW.

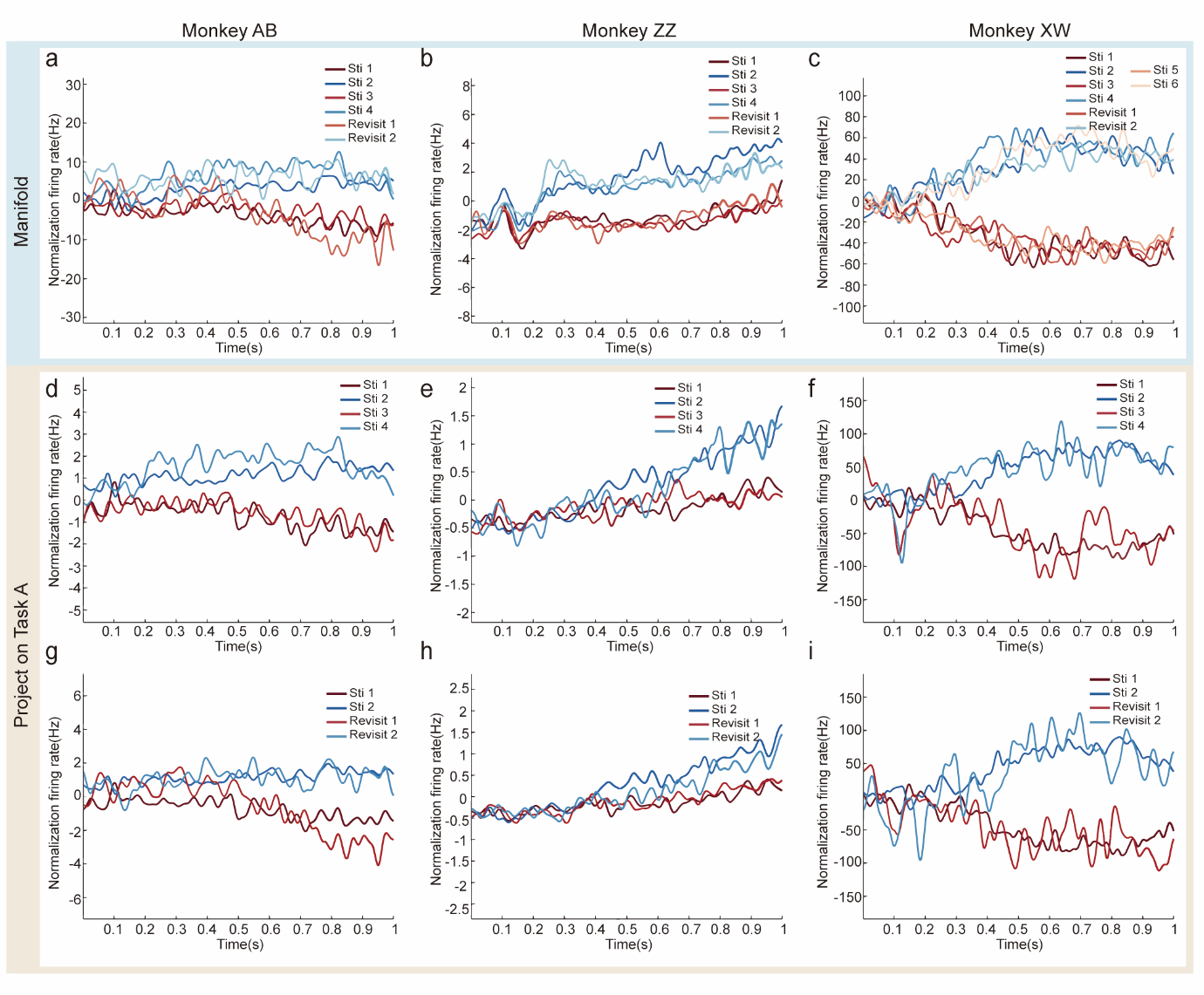

Supplementary Fig. 4. Decision-related subspace manifolds in three monkeys. (a, b, c) Neural activities from different tasks, including Tasks A, B/C, and Revisit A, was projected onto a common decision subspace for each of the three monkeys. (d, e, f, g, h, i) To verify whether the manifold formed in Task A was reused in subsequent tasks: (d, e, f) Neural activity from Task B was projected onto the same decision subspace of Task A; (g, h, i) Neural activity from the Revisit A task was projected onto the same decision subspace of Task A. Sti, stimulus; Revisit, Revisit A.

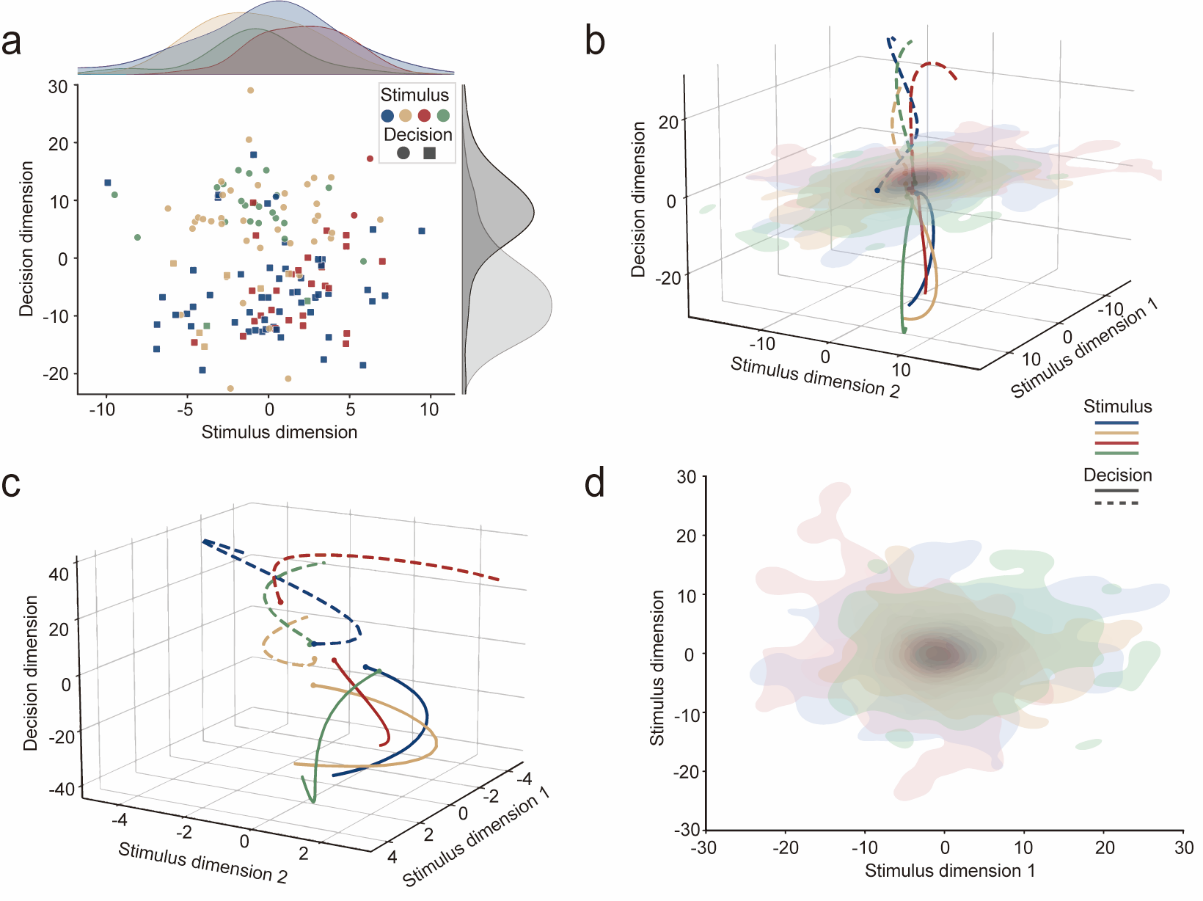

Supplementary Fig. 5. Orthogonal relationship between decision subspace and stimulus subspace: an example from monkey ZZ on one day. (a) Distribution of different trials on the stimulus-decision plane; (b) The neural dynamics trajectory in the three-dimensional space of the decision and stimulus subspaces; (c) Neural trajectories of motor decision dynamics in three-dimensional space; (d) Projection of neural activity in the visual stimulus subspace.
